# A burst of regulatory and protein innovation at the origin of placental mammals drove the emergence of placenta and underpins divergent early pregnancy strategies in modern mammals

**DOI:** 10.1101/2021.07.22.453388

**Authors:** Alysha S. Taylor, Haidee Tinning, Vladimir Ovchinnikov, William Smith, Anna L. Pullinger, Ruth A. Sutton, Bede Constantinides, Dapeng Wang, Niamh Forde, Mary J O’Connell

## Abstract

The origin of live birth in mammals ∼148 million years ago was a dramatic shift in reproductive strategy, yet the molecular changes that established mammal viviparity are largely unknown. Although progesterone receptor signalling predates the origin of mammals andis highly conserved in, and critical for, successful mammal pregnancy, it alone cannot explain the origin and subsequent diversity of implantation strategies throughout the placental mammal radiation. MiRNAs are known to be flexible and dynamic regulators with a well established role in the pathophysiology of mammal placenta. We propose that a dynamic core microRNA (miRNA) network originated early in placental mammal evolution, responds to conserved mammal pregnancy cues (*e*.*g*. progesterone), and facilitates species-specific responses. Here we identify 13 miRNAs that arose at the origin of placental mammals and were subsequently retained in all descendent lineages. The expression of these 13 miRNAs in response to early pregnancy molecules is regulated in a species-specific manner in endometrial epithelia of species with extremes of implantation strategies. Furthermore, these 13 miRNAs preferentially target 84 proteins under positive selective pressure on ancestral eutherian. Discovery of this core “live-birth” toolkit and specifically adapted proteins helps explain the origin and evolution of the placenta in mammals.

---

Successful pregnancy in eutheria is contingent on a developmentally competent embryo, appropriate endometrial function, and on formation of the placenta and on the molecular cross-talk across these components ^1,2^. Yet placental organ morphology, and more generally, the underlying regulation of successful pregnancy, varies across mammals. Protein coding alterations along with innovations in regulatory networks drive the origin and evolution of novel traits ^3^. Moreover, bursts of evolution of new microRNAs (miRNAs) are known to be associated with morphological innovation ^4–6^, and miRNAs are known to regulate placental function in both normal and pathophysiological conditions ^7,8^. We have uncovered a core placental toolkit in mammals that evolved through a synergy of molecular events involving the emergence of new miRNAs combined with adaptive amino acid changes in key proteins. We propose that this conserved core toolkit contributes to the diversity of implantation strategies and pregnancy morphologies observed in modern eutheria (Figure 1).

**Figure 1.**
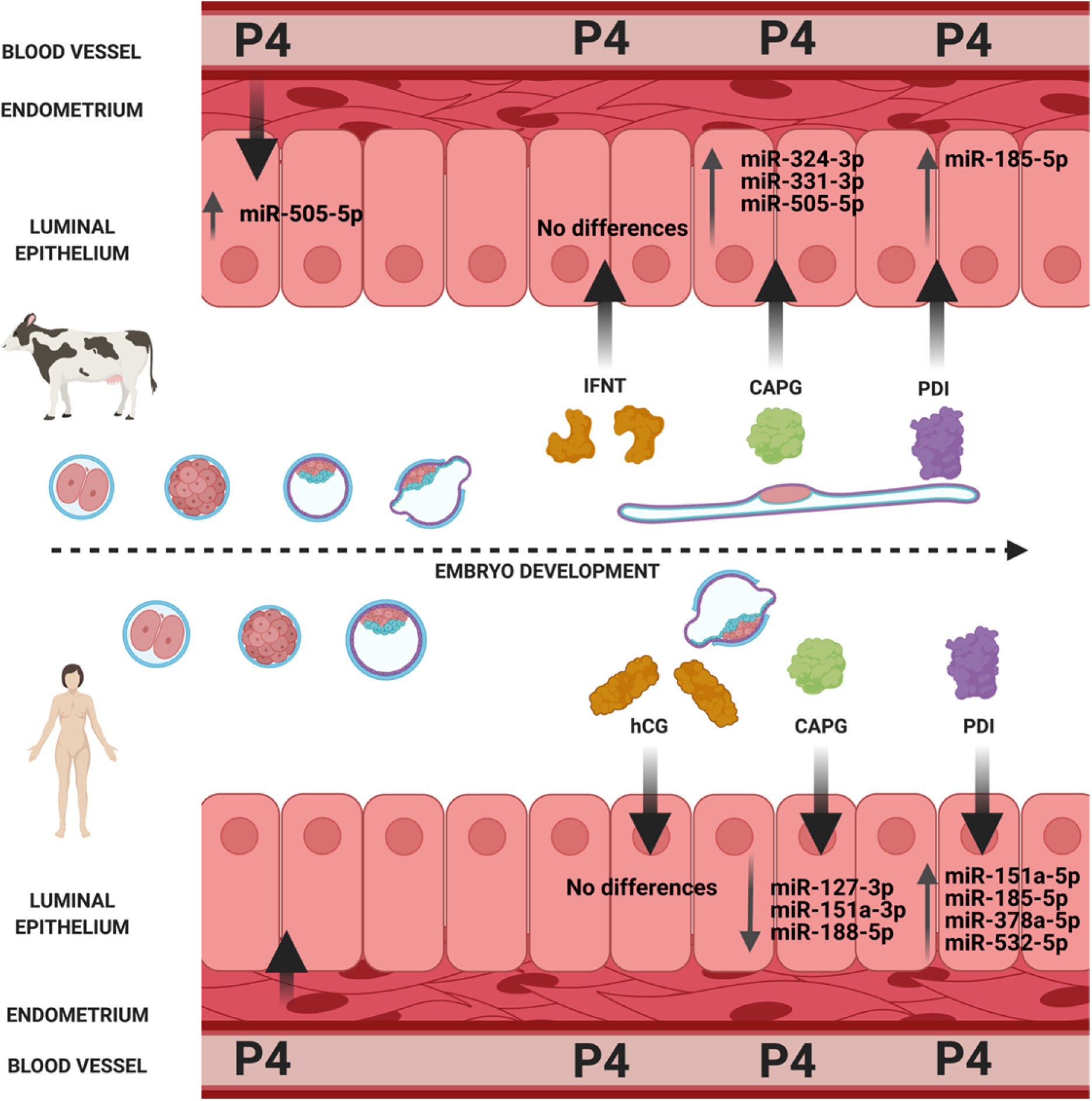
Regulation of core miRNA toolkit by early pregnancy markers. The endometrial epithelium of bovine (superficial implantation strategy) (top panel) or human (invasive implantation strategy) (bottom panel) are regulated by molecules important for endometrial function in early pregnancy in eutheria (P4: progesterone; hCG: human chorionic gonadotrophin; Interferon Tau: IFNT; Macrophage capping protein: CAPG, and protein disulfide isomerase: PDI).

## A core miRNA toolkit of 13 miRNAs arose at the origin of therian and eutherian mammals and were never subsequently lost

Because significant miRNA family expansions have been found to correlate with major transitions in animal evolution^4–6^, we interrogated MirGeneDB^9^ and identified 112 miRNA families that originated on either the therian (6 miRNA families) or eutherian stem lineage (106 miRNA families). Six of these miRNA families emerged on the older therian stem lineage (mir-340, mir-483, mir-671, mir-675, mir-1251, and mir-3613), of which only mir-340 is phylogenetically conserved in both Theria and Eutheria (Figure 2a). Given that mir-671 is present in tasmanian devil and in all Eutheria sampled, we classified mir-671 as a ‘therian stem lineage miRNA’. A total of 106 miRNAs emerged at the origin of eutheria, and 11 of these remained conserved in all extant eutheria sampled (Figure 2a). In total, this yielded 13 miRNAs that originated on stem mammal lineages and were subsequently retained in all descendent lineages, henceforth referred to as “the core miRNA toolkit”. Expression and function of some of the core miRNA toolkit has been demonstrated for normal placenta (*e*.*g*. mir-433^10^, mir-28^11^, mir-378^12^), and endometrium (*e*.*g*. mir-505^13^ and mir-542^14^). Others have been implicated in pathophysiological pregnancies, *e*.*g*. mir-185, mir-188, mir-423 and mir-542^15–17^ have been implicated in preeclampsia, mir-127^10^ has been associated with placentomegaly, mir-324 is associated with LGA pregnancies^18^, mir-331 is associated with placenta from intra-amniotic infection^19^, and mir-505 can be associated with preterm birth^20^. Collectively all 13 members of the core miRNA toolkit have been implicated in critical roles in placental mammal pregnancies.

**Figure 2:**
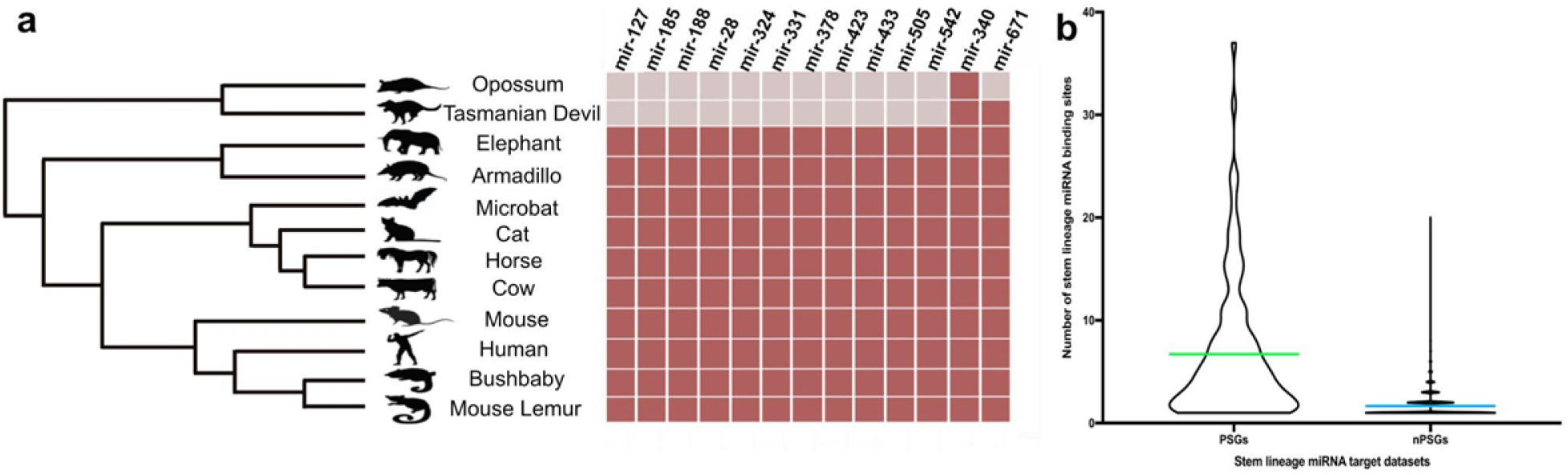
Phylogenetic distribution of the miRNA families and their levels of targeting in positively selected genes. **(a)** Phylogenetic distribution of miRNA families specific to therian and eutherian mammals. The mammal phylogeny displaying the species sampled in our analysis. The corresponding matrix shows the presence (dark red) or absence (pale grey) of miRNA families across the species sampled. **(b)** Violin plot comparing the number of target miRNA binding sites from the core miRNA toolkit per transcript in the 84 genes that underwent positive selection on the stem Eutherian lineage (PSGs) (left) compared to sets of randomly sampled targets of the miRNAs that do not have signatures of positive selection on the stem Eutherian lineage (nPSG) (right). For each of the 84 PSGs, the mean number of predicted miRNA binding sites was determined for each target transcript in human (depicted in green). This was compared to the mean number of binding sites for each of the nPSGs (depicted in blue). The mean number of binding sites was determined to be significantly different between the two datasets when p≤0.05, two-sample t-test with unequal variance.

Targets were identified in the human genome for the core miRNA toolkit using TargetScan^21^. The predicted functions of the targets of the core miRNA toolkit include reproductive functions (55 target transcripts), metabolic process (1476 target transcripts), and Biological regulation (1269 target transcripts) (Supplementary Figure S1b). More specifically, some of the targets were implicated in pathways associated with INPP5E regulation, Neurophilin interactions, and VEGF interactions with its receptor (VEGR), each involved in the process of angiogenesis. TGF-β signalling and p53 regulation are amongst the predicted targets and are implicated in cell proliferation (Supplementary Figure S3). Both angiogenesis and proliferation are required for successful pregnancy. The syncytins are a family of endogenous retrovirus-derived protein coding regions that were domesticated in mammals and are essential for promoting placental formation^22^ and 4/13 of the core miRNA toolkit (miR-185-3p, miR-188-3p, miR-423-5p and miR-433-3p) are predicted to target the syncytins with at least 7-mer binding. In the case of miR-423-5p, there are two predicted target sites for *syncytin-1*, one of which has a site overlapping with that of the miR-185-3p binding site, indicating dynamic/competitive binding between these miRNAs and their *syncytin-1* target. The predicted targets of the core miRNA toolkit also include 130 gene families that have been proposed to have emerged on the eutherian stem lineage^23^. In addition, the targets included genes that evolved endometrial expression on the stem eutherian lineage, and that are hypothesized to have assisted in the remodelling of the uterine landscape during the evolution of the mammal placenta^1^.

### Evidence of positive selection on the stem eutherian lineage

Protein coding alterations (such as the birth of new genes and gene families, gene loss, co-option, and selective pressure variation) along with innovations in regulatory networks drive the development of novel traits^3^. A number of cases of positively selected amino acids (indicative of protein functional shift) are known to have had a direct role in endometrial function, *e*.*g*. the galectin family of proteins involved in immune modification at the maternal-fetal interface^24^. We looked for evidence of adaptive evolution in single gene orthologous families (SGOs) on the stem eutherian lineage. We focussed on SGOs to optimise our ability to assign function to orthologs and to accurately trace evolutionary histories. We chose a total of ten Eutheria that demonstrate the greatest range of diversity in placental morphology, plus four outgroup species, one from each of the *Monotremata, Marsupialia, Aves* and *Teleostei* (Supplementary Table S1). Annotated gene families were taken from Ensembl 90^25^. Following our filtering regime we extracted a total of 1,437 SGOs. Applying codon models of evolution to these SGOs, we identified signatures of positive selection on amino acid residues on the stem eutherian lineage in 237 SGOs. The functions of these genes are predominantly cellular processes, metabolic processes, and biological regulation (Supplementary Figure S2). Out of these 237 positively selected SGOs, 115 contained positively selected amino acid residues that were subsequently unaltered in all descendent lineages. The 115 SGOs are functionally enriched for chromosomal maintenance, telomere activity, p53 signalling, cell cycle, and the inflammatory immune response - activities that have been associated with the formation of the placental tissue in pregnancy^26–28^. We then studied whether there is a significant association between the core miRNA toolkit and stem-lineage positively selected proteins.

### The core miRNA toolkit preferentially targets genes under positive selection in the stem eutherian lineage

Synergy between regulatory and protein coding innovations often drives substantial phenotypic novelty^2,3,29^. Therefore, to test the hypothesis that innovation both at the level of regulation and of protein coding change underpinned the origin of placentation in mammals, we performed a simulation study on the targets of the 13 stem lineage miRNAs. Out of 115 SGOs with evidence of positive selection on the ancestral eutherian lineage, 84 were found to be significantly enriched for binding sites for the 13 members of the core miRNA toolkit (p=1.35618e-11), with a mean of 6.66 binding sites per transcript (median=4.0) (Figure 2b). We determined if the number of binding sites in this subset of 84 positively selected SGOs was significantly different than one would expect by random chance. We estimated the number of binding sites per transcript for the core miRNA toolkit in 100 randomly sampled gene sets and found it is significantly lower, with a mean=1.64 (median=1.0). This suggests that the 13 miRNAs in the core miRNA toolkit preferentially target the positively selected SGOs (6.66 binding sites compared to 1.64) (Figure 2b). Combined, this indicates that a co-evolutionary process arose in a short time in early mammal evolution that resulted in altered protein function, as well as a new miRNA-mediated regulatory network.

The functions of the 84 positively selected SGOs targeted by the core miRNA toolkit broadly fall within the categories of cell cycle, DNA damage & DNA metabolic processes, and regulation of hair cycle & hair cycle – a uniquely mammal characteristic (Supplementary Figure S3). 21/84 SGOs are significantly more likely to interact with one another (p<0.05) in comparison to any other gene in the genome (Supplementary Figure S2b). Of course there is significant variation found in modern mammals in other facets such as telomere biology, cancer incidence, body mass and maximum lifespan^30–33^, therefore innovation at this node was not expected to be entirely skewed to pregnancy (Supplementary Table S2)

### Species-specific regulation of the core miRNA toolkit by key early pregnancy molecules in species with different implantation strategies

Implantation in eutherian mammals displays wide variation in both embryological morphology and bi-lateral signalling between the embryo and maternal endometrium, and degree of invasiveness (invasive in human, superficial in bovine). This variation is governed, in part, by conserved signalling pathways, *e*.*g*. the sustained actions of the hormone progesterone (P4)^34^, but also diverse molecular cues such as the maternal recognition of pregnancy signal, *e*.*g*. chorionic gonadotrophin (hCG) in human and interferon (IFNT) in bovine^35,36^. We asked the question what molecules, that are involved in bi-lateral communication between the embryo and endometrium, regulate the core mammal miRNA toolkit and if they are regulated in a species-specific manner. We cultured endometrial epithelial cells from human and bovine^37^ and exposed them for 24 hr to recombinant forms of conserved (P4, CAPG, and PDI) and diverse (IFNT and hCG) molecular cues important for early pregnancy success in placental mammals. CAPG and PDI have recently been identified as produced by the bovine conceptus during pregnancy recognition and are highly conserved (in terms of sequence identity and phylogenetic distribution) across placental mammals^37,38^. We then examined the expression of the 13 stem lineage miRNAs in these cells using an LNA-based approach. Treatment of bovine endometrial epithelial cells with 10μg/mL of P4 during the early luteal phase resulted in increased expression of miR-505-5p (Supplementary Figure S4). In summary, treatment with the evolutionarily conserved early pregnancy proteins (P4, CAPG and PDI) in the endometrial epithelial cells of bovine and/or human resulted in a change in expression of 11/13 of the stem lineage miRNAs (Figure 1).

Intriguingly, addition of recombinant bovine forms of bCAPG and bPDI proteins to bovine or human endometrial epithelial cells altered expression of selected miRNAs in a species-specific manner (Figure 3&4 respectively). Treatment of bovine cells with recombinant bCAPG resulted in increased expression of miR-331-3p, miR-324-5p and miR-505-5p in endometrial epithelial cells (p<0.05). Whereas in human endometrial epithelial cells treated with 1000ng/μl bCAPG, the expression of miR-127-3p, miR-151a-3p (a paralog of mir-28 originating on the Eutherian stem lineage), and miR-188-5p showed a significant decrease in expression compared to vehicle control (p<0.05: Figure 3). Treatment with recombinant bPDI decreased expression of miR-185-5p in bovine epithelial cells (P<0.05). In human Ishikawa immortalised endometrial epithelial cells treated with 1000ng/μl bPDI the expression of miR-151a-5p, miR-185-5p, miR-378a-3p and miR-532-5p (a paralogue of mir-188 originating on the Eutherian stem lineage) were significantly decreased compared to vehicle control (Figure 4).

**Figure 3.**
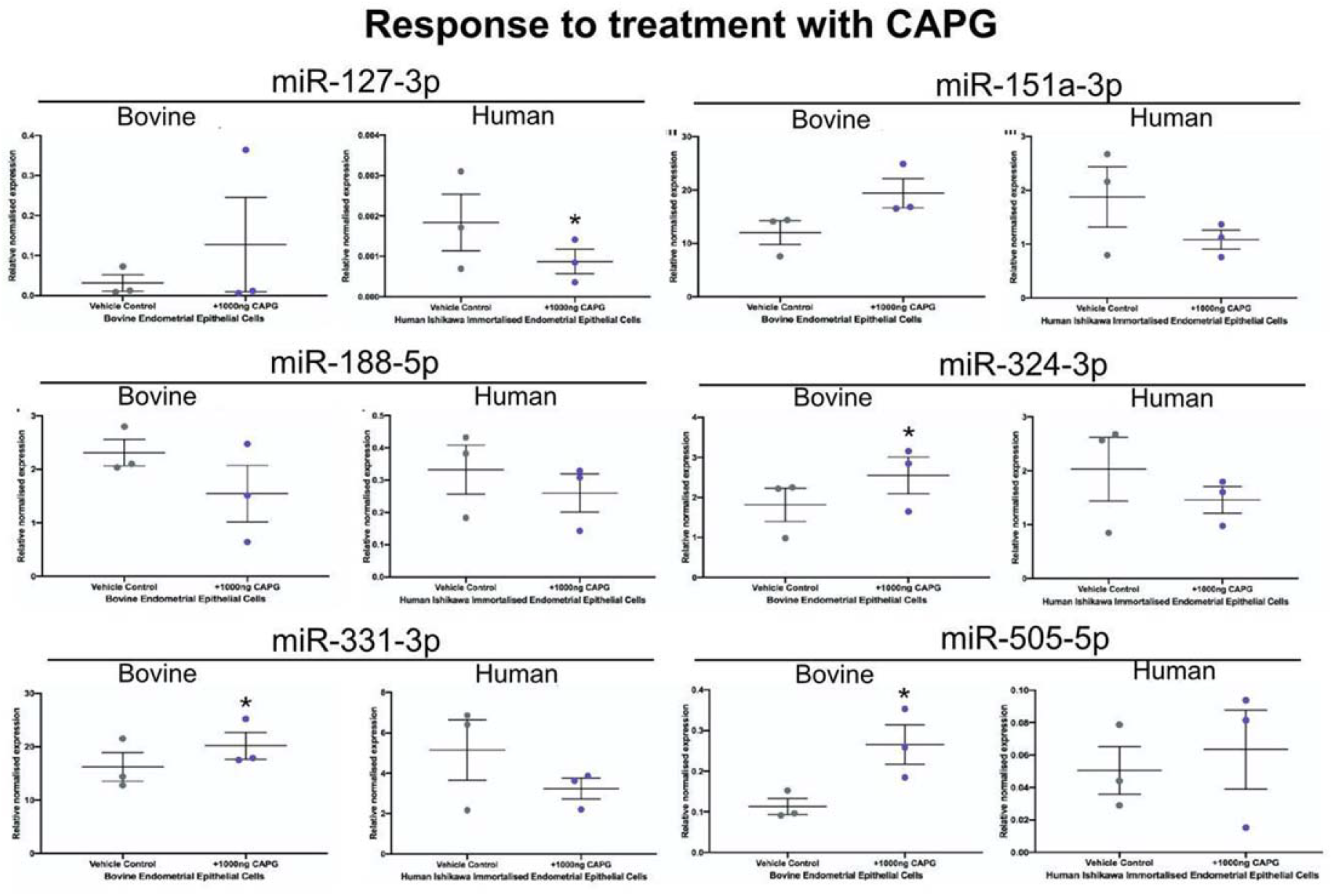
Regulation of stem lineage miRNAs in bovine and human endometrial epithelial cells treated with bCAPG. Expression of stem lineage miRNAs miR-127-3p, miR-151a-3p, miR-188-5p, miR-324-3p, miR-331-3p, and miR-505-5p in either bovine (left hand side of each pair) or human (right hand side of each pair) endometrial epithelial cells. Primary bovine endometrial epithelial cells were treated with vehicle control (grey circle), or 1000ng/μl bCAPG (purple circle) for 24 hours. Human Ishikawa immortalized endometrial epithelial cells were treated with vehicle control (grey circle), or 1000ng/μl bCAPG (purple circle) for 24 hours. Significant differences in miRNA expression values determined when p≤0.05 are depicted by an asterisk (*).

**Figure 4:**
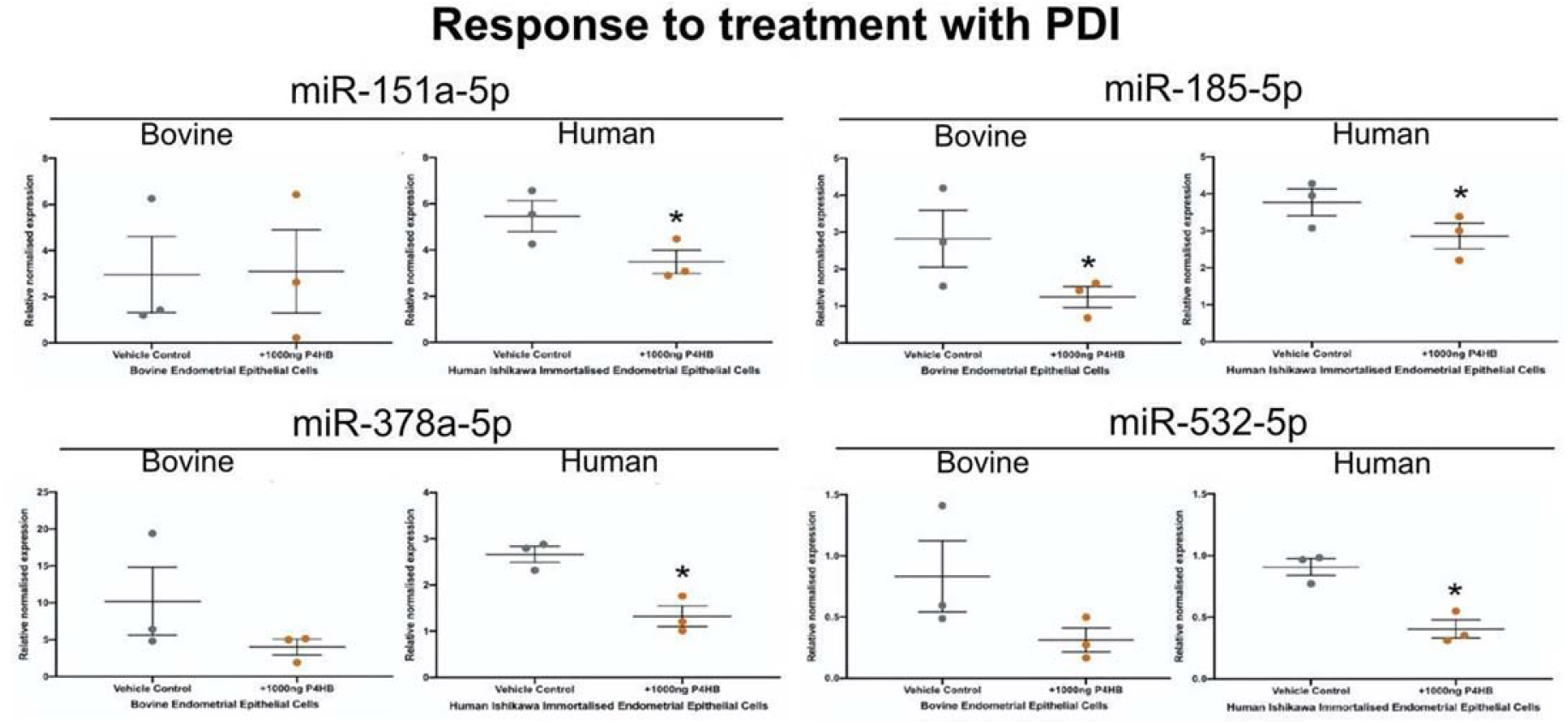
Regulation of stem lineage miRNAs in bovine and human endometrial epithelial cells treated with bPDI. Expression of stem lineage miRNAs miR-127-3p, miR-151a-3p, miR-188-5p, miR-324-3p, miR-331-3p, and miR-505-5p in either bovine (left hand side of each pair) or human (right hand side of each pair) endometrial epithelial cells. Primary bovine endometrial epithelial cells were treated with vehicle control (grey circle), or 1000ng/μl bPDI (orange circle) for 24 hours. Human Ishikawa immortalized endometrial epithelial cells were treated with vehicle control (grey circle), or 1000ng/μl bPDI (orange circle) for 24 hours. Significant differences in miRNA expression values determined when p≤0.05 are depicted by an asterisk (*).

In contrast, addition of the species specific pregnancy recognition signals (IFNT in bovine: hCG in human) to receptive endometrial epithelial cells altered expression of one member of the core miRNA toolkit (Supplementary Figures S5 and S6 for IFNT and hCG data respectively). These data demonstrate that the expression of the core miRNA toolkit is not altered by the species-specific pregnancy recognition signals (IFNT and hCG).

Human and bovine represent two distinct implantation strategies for mammals and these lineages last shared a common ancestor some ∼92 million years ago, representing ∼184 million years of independent evolution ^39,40^. None of the 13 stem lineage miRNAs are regulated by the species-specific pregnancy recognition signals (IFNT and hCG), but the expression of the core miRNA toolkit is modified by proteins that are highly conserved amongst placental mammals (CAPG and PDI). Combined, our results show that the preferential targeting of the core mammal miRNA toolkit, and protein functional shift were essential to the establishment of mammalian implantation, and that subsequent diversification of this network facilitated the range of implantation strategies observed today.

In summary, this work identifies a core regulatory network that drove a major transition - the origin of live birth in mammals. With the core now defined, future work can focus on the accessory elements that drove the subsequent diversification of mammal placenta.

## ACKNOWLEDGEMENTS

We would like to thank Dr Fuller Bazer from Texas A&M for the kind gift of recombinant ovine IFNT. We would like to acknowledge the assistance of Stefania Mountevedi in helping with the bovine cell isolation and culture. Recombinant proteins CAPG and PDI were generated by the Protein and Proteome Analysis core (NUPPA), Newcastle University, Newcastle upon Tyne, UK. This work was funded by University of Leeds under the LARS PhD scholarship (AST) and Maternity leave support scheme to MOC for BC, and by the University of Nottingham postdoctoral fellowship funding to MOC for VO. This work was supported by N8 agri-food pump priming, QR GCRF, as well as BBSRC grant number BB/R017522/1 to NF. This work was undertaken on ARC3, part of the High-Performance Computing facilities at the University of Leeds, UK. Figure 1 was created using Biorender.

## CONTRIBUTIONS

NF and MJOC conceived of the study and designed the experiments. AST, BC, VO and MJOC carried out all evolutionary analyses and DW carried out statistical tests. NF, AST, HT, WS, ALP, and RAS undertook all molecular analyses. NF and MJOC drafted the manuscript, all authors contributed to revising, reading, and critiquing the manuscript.

## Supplementary Figures and Tables

**Figure S1:**
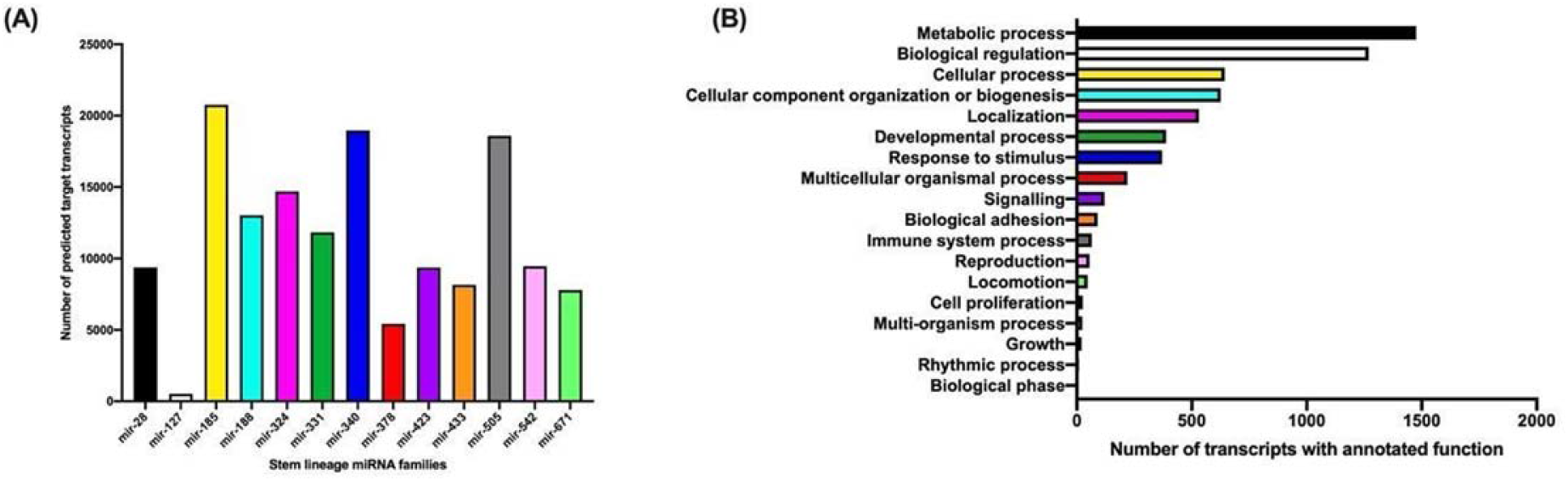
Number of predicted targets for each of the 13 stem lineage miRNA family and the functional annotation of the target genes using Panther DB. **(A)** Targets were predicted for each of the 13 stem lineage miRNAs using TargetScan70 (Agarwal et al., 2015). TargetScan output was filtered for targets with8mer-A1, 7mer-m8 and 7mer-A1 complementary binding to the seed region. **(B)** Functional annotation of predicted targets of the 13 stem lineage miRNAs. Filtered stem lineage miRNA target transcripts were analysed for functional enrichment using PANTHERv.14 (Muruganujan et al., 2018), where PANTHERv.14 found functional annotations to be significantly enriched when p≤0.05.

**Figure S2:**
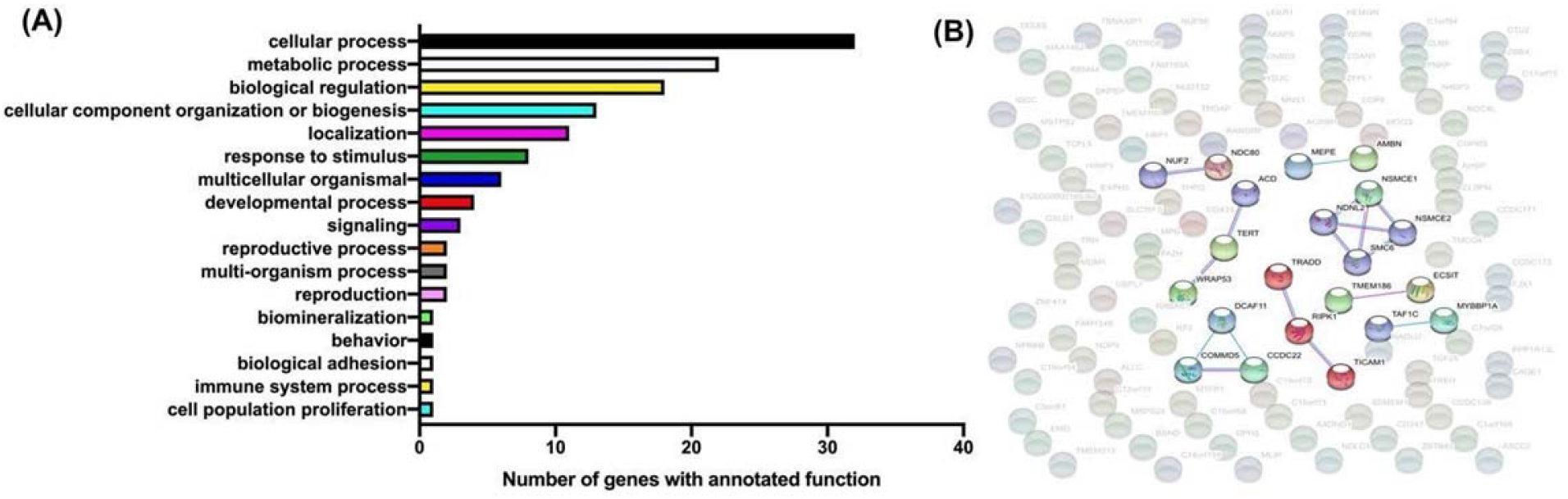
Broad functions of the genes with signatures of positive selection and sample of their interactions. **(A)** Functional annotation using Gene Ontology Biological Process terms of 115 SGOs that underwent positive selection on the stem eutherian lineage and where the amino acid substitution was fixed on all extant Eutheria tested. The absolute number (out of 115) of positively selected genes in a given category are shown in the X axis and the functional annotations on the Y-axis. **(B)** String interaction network of the same set of 115 genes. Network has 106 total nodes and 8 edges (expected edges =4). Background nodes, with no high confidence interactions from experimental and database sources are faded. Nodes with high confidence interactions from experimentally determined (pink lines) or curated database (blue lines) sources are depicted in colour. The network was found to be significantly enriched for gene-gene interactions (p=0.0468). Average node degree is 0.151, with an average local clustering coefficient of 0.104. Minimum interaction score is 0.700.

**Figure S3:**
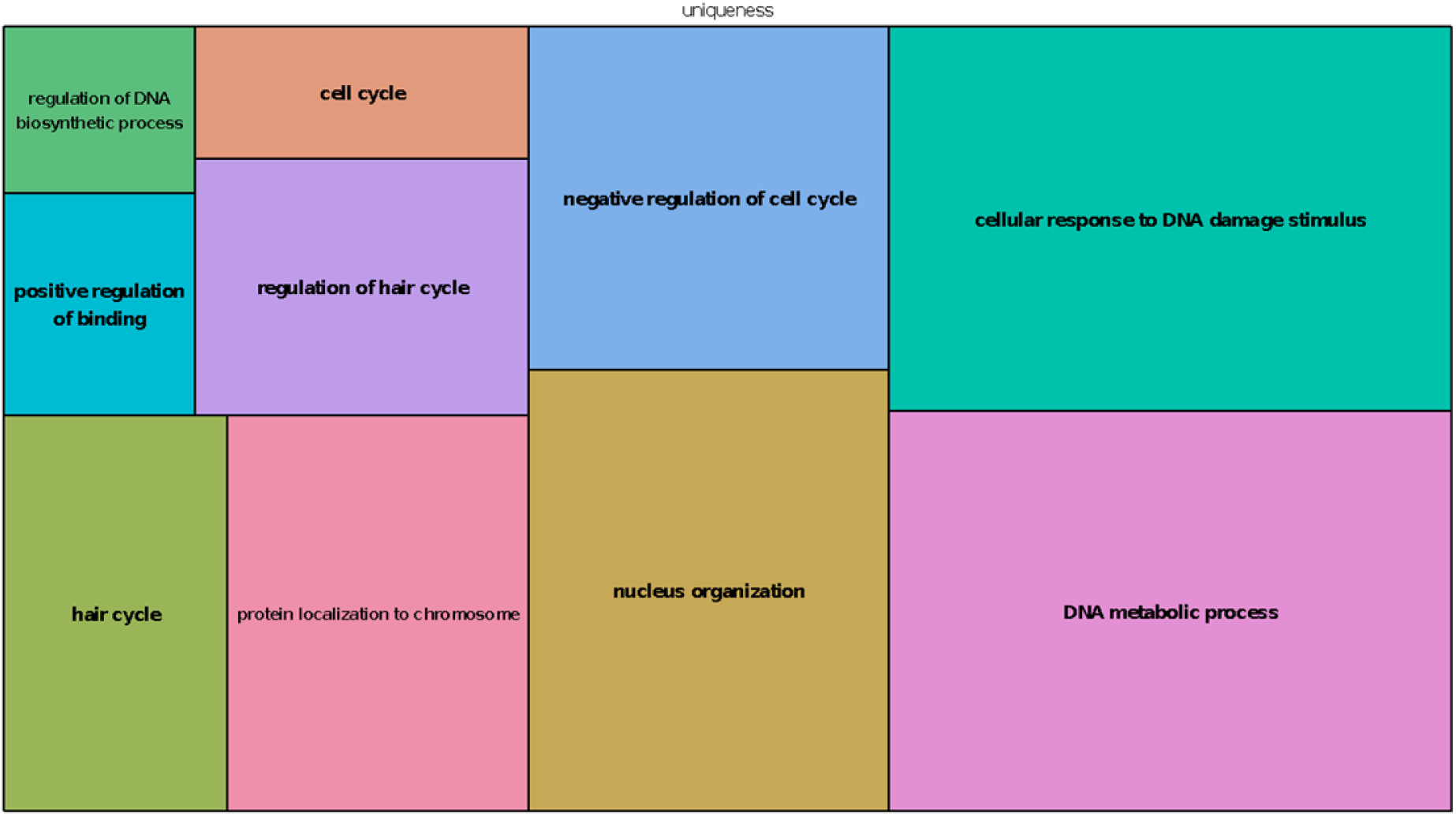
A standard TreeMap from REVIGO displaying the GO biological process terms present in the 84 PSGs that were predicted targets of the 13 stem lineage miRNAs. Rectangle size represents semantic uniqueness of GO term, defined by REVIGO as the negative of average similarity to all other terms present in human.

**Figure S4:**
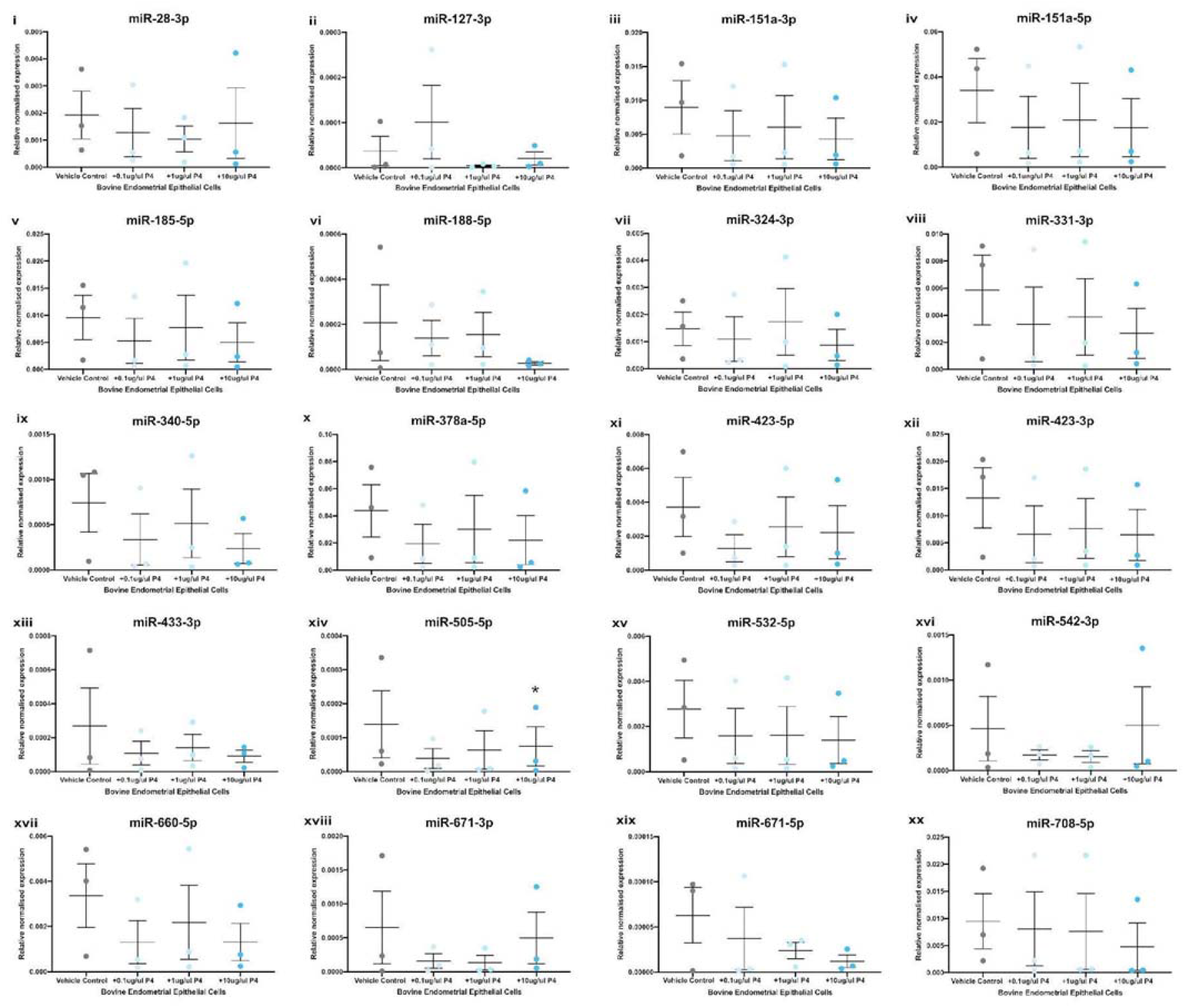
Expression of stem lineage miRNAs in bovine endometrial epithelial cells treated with P4. Expression of stem lineage miRNA (i) miR-28-3p, (ii) miR-127-3p, (iii) miR-151a-3p, (iv) miR-151a-5p, (v) miR-185-5p, (vi) miR-188-5p, (vii) miR-324-5p, (viii) miR-331-3p, (ix) miR-340-5p, (x) miR-378a-5p, (xi) miR-423-3p, (xii) miR-423-5p, (xiii) miR-433-3p, (xiv) miR-505-5p, (xv) miR-532-5p, (xvi) miR-542-3p, (xvii) miR-660-5p, (xviii) miR-671-3p, (xix) miR-671-5p and (xx) miR-708-5p in bovine endometrial epithelial cells treated with vehicle control (grey circle), 0.1μg/mL (light blue), 1.0μg/mL (medium blue) or 10μg/mL P4 (dark blue circle) for 24 hours. Significant differences in miRNA expression values determined when p≤0.05 are depicted by an asterisk (*).

**Figure S5:**
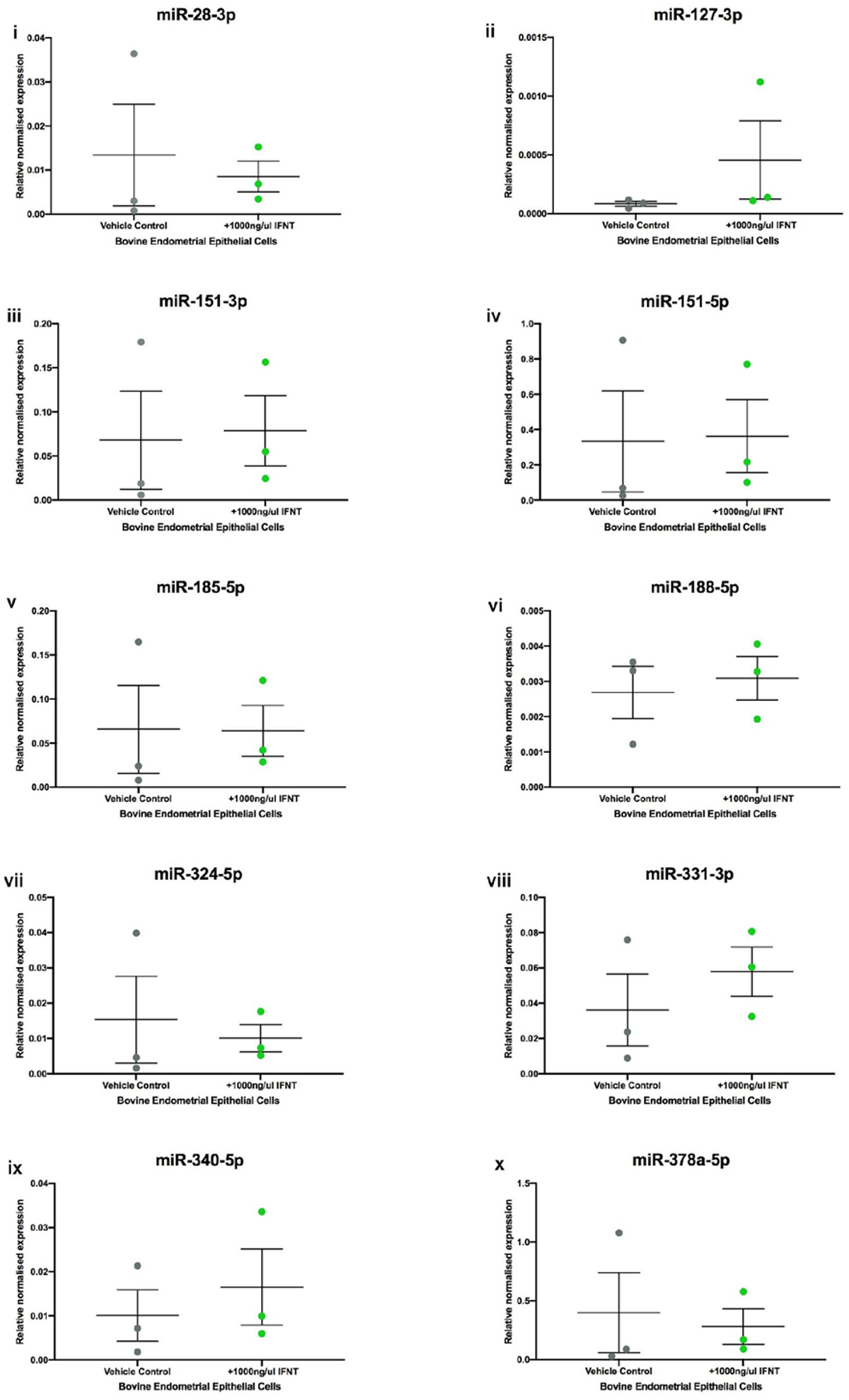
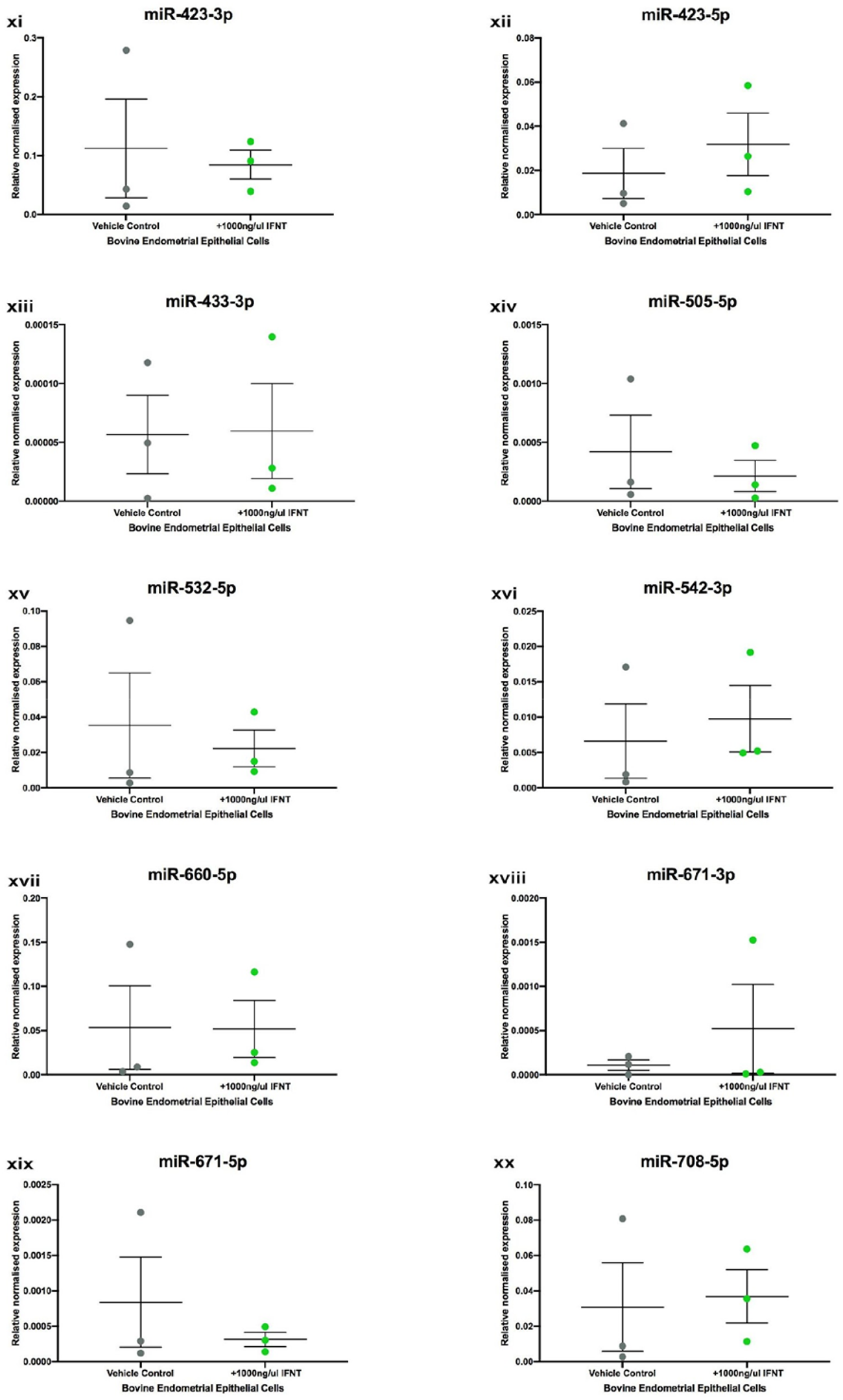
Expression of stem lineage miRNAs in bovine endometrial explants treated with recombinant oIFNT. Expression of stem lineage miRNA (i) miR-28-3p, (ii) miR-127-3p, (iii) miR-151a-3p, (iv) miR-151a-5p, (v) miR-185-5p, (vi) miR-188-5p, (vii) miR-324-5p, (viii) miR-331-3p, (ix) miR-340-5p, (x) miR-378a-5p, (xi) miR-423-3p, (xii) miR-423-5p, (xiii) miR-433-3p, (xiv) miR-505-5p, (xv) miR-532-5p, (xvi) miR-542-3p, (xvii) miR-660-5p, (xviii) miR-671-3p, (xix) miR-671-5p and (xx) miR-708-5p in bovine endometrial explants treated with vehicle control (grey circle), or 1000ng/μl oIFNT (green circle) for 24 hours. Significant differences in miRNA expression values determined when p<0.05 are depicted by an asterisk (*).

**Figure S6:**
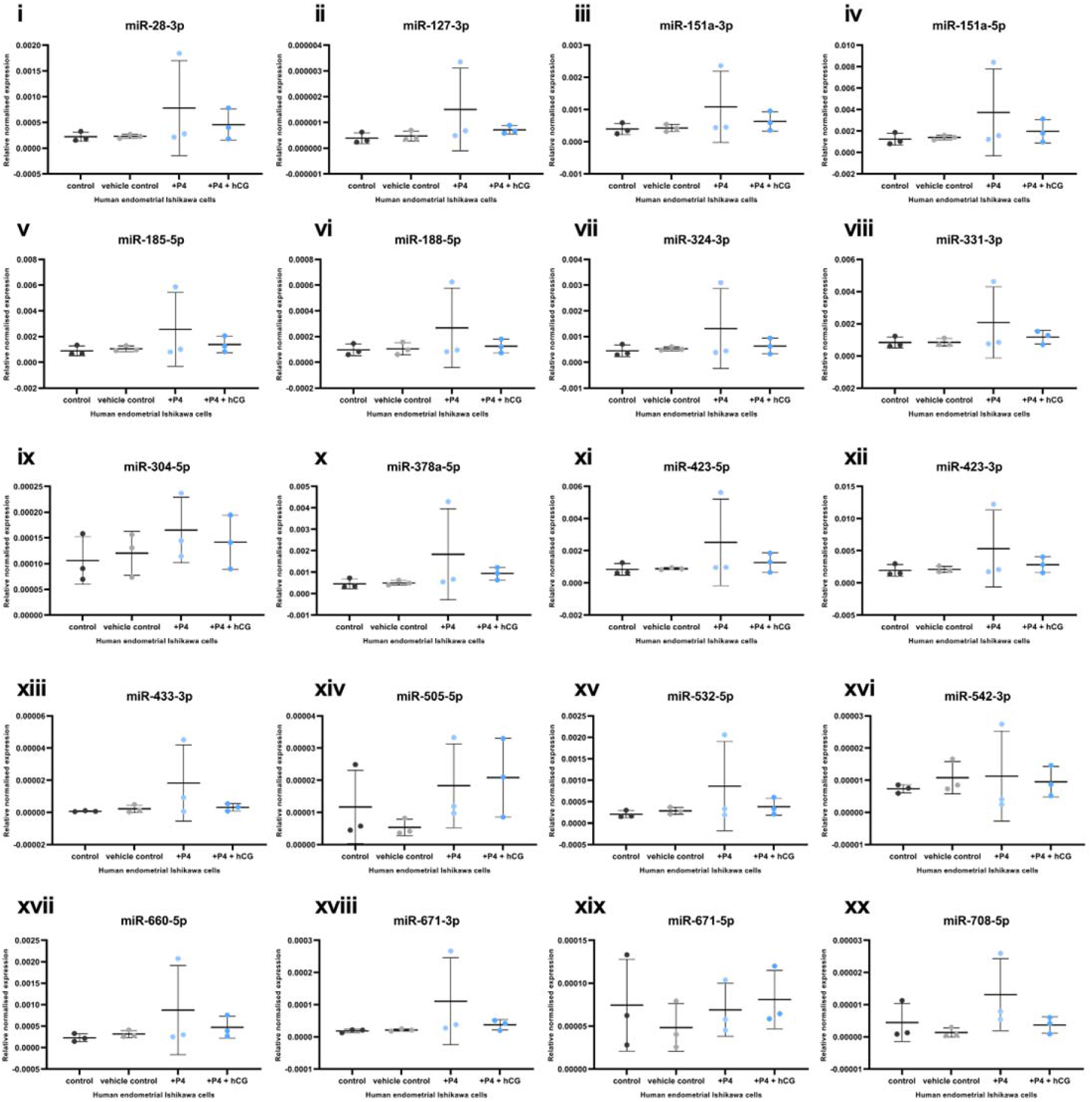
Expression of stem lineage miRNAs in human endometrial epithelial cells treated with hCG. Expression of stem lineage miRNA (i) miR-28-3p, (ii) miR-127-3p, (iii) miR-151a-3p, (iv) miR-151a-5p, (v) miR-185-5p, (vi) miR-188-5p, (vii) miR-324-5p, (viii) miR-331-3p, (ix) miR-340-5p, (x) miR-378a-5p, (xi) miR-423-3p, (xii) miR-423-5p, (xiii) miR-433-3p, (xiv) miR-505-5p, (xv) miR-532-5p, (xvi) miR-542-3p, (xvii) miR-660-5p, (xviii) miR-671-3p, (xix) miR-671-5p and (xx) miR-708-5p in human Ishikawa immortalized endometrial epithelial cells were treated with control (dark grey circle), vehicle control (light grey circle), or P4 (light blue circle), or P4+hCG (darker blue circles) for 24 hours. No differences in miRNA expression were determined (p>0.05).

**Supplementary Table S1:** Vertebrate Species sampled, genome version, coverage and completion level. Set of 14 species sampled for the selective pressure analysis. Using Ensembl 92 (Yates et al., 2016), a dataset of genomes representative of (i) vertebrate outgroup clades, or (ii) variations in placental morphology in metatherian and eutherian mammals. For each clade, genomes were chosen based on genome coverage for downstream homology searching. ‘Clade’ refers to the taxonomic group of each included species. ‘Species’ denotes the included species, by their common name. ‘Version’ denotes the genome assembly version included in this analysis. ‘Genome Quality’ refers to the coverage and assembly level of each species included in this analysis.

| Clade | Species | Version | Genome Quality |
| --- | --- | --- | --- |
| **Fish** | Zebrafish | GRCz11 | Full Genome, Chromosome Level |
|  | Chicken | Gallus_gallus-5.0 | 70X Coverage, Chromosome Level |
| **Monotremes** | Platypus | OANA5 | 6X Coverage, Chromosome Level |
| **Metatheria** | Opossum | monDom5 | 7.33X Coverage, Chromosome Level |
| **Eutheria** | Elephant | Loxafr3.0 | 7X Coverage, Scaffold Level |
|  | Armadillo | Dasnov3.0 | 6X Coverage, Scaffold Level |
|  | Mouse | GRCm38.p6 | High Quality Reference Assembly |
|  | Human | GRCh38.p12 | High Quality Reference Assembly |
|  | Mouse Lemur | Mmur_3.0 | 221.6X Coverage, Chromosome Level |
|  | Bushbaby | OtoGar3 | 137X Coverage, Scaffold Level |
|  | Cat | Felis_catus_9.0 | 72X Coverage, Chromosome Level |
|  | Microbat | Myoluc2.0 | 7X Coverage, Scaffold Level |
|  | Horse | Equ Cab 2 | 6.79X Coverage, Chromosome Level |
|  | Cow | UMD3.1 | 9X Coverage, Chromosome Level |

**Supplementary Table S2:** Example of three functionally related proteins under positive selection on stem eutherian lineage. The panther family ID and common gene names are provided for a set of 3 proteins extracted to illustrate the cases of positive selection identified. The LnL value associated with the fit of the codeml model (PAML) to the data are provided as are the associated proportion of sites (p2) that have the corresponding ω2 (or Dn/Ds ratio). The sites estimated to be positively selected are given in the final column, these are numbered as per aligned codon position.

| Family | Gene Name | LnL | $p2$ | $\omega2$ | Positions in the alignment predicted to be positively selected. |
| --- | --- | --- | --- | --- | --- |
| PTHR13211 | WRAP53 | -10149.383 | 0.04821 | 37.80743 | 72, 88, 114, 225, 253, 260, 261, 331, 423, 480, 493, 506, 510, 513, 522, 528 |
| PTHR12066 | TERT | -36003.667 | 0.02302 | 954.86770 | 121, 122, 140, 166, 197, 508, 878, 1020, 1110 |
| PTHR14487 | ACD | -15296.106 | 0.10315 | 30.62227 | 138, 143, 148, 178, 185, 200, 202, 206, 208, 209, 214, 269, 278, 296, 298, 314, 347, 363, 368, 379, 390, 394, 395, 412, 414, 450, 470, 474, 524, 530, 532, 533, 545, 548, 561, 571, 575, 586, 685, 698 |

